# The virulence regulator *bvgS* is required for nutrient-induced filamentation in *Bordetella avium*

**DOI:** 10.1101/2025.01.19.633786

**Authors:** Niklas G Perslow, Robert J Luallen

## Abstract

Bacteria can change morphology in response to stressors and changes in their environment, including infection of a host. We previously identified the bacterial species, *Bordetella atropi*, which uses nutrient-induced filamentation as a novel mechanism for cell-to-cell spreading in the intestinal epithelial cells of a nematode host. To further investigate the conservation of nutrient-induced filamentation in Bordetellae, we utilized the turkey-infecting species *Bordetella avium* which filaments in vitro when switched from a standard growth media to an enriched media. We conducted a selection-based filamentation screen with *B. avium* and isolated two independent non-filamentous mutants that failed to filament in highly enriched media. These mutants contained different alleles in *bvgS*, the sensor in the two-component master virulence regulator (BvgAS) conserved across the Bordetella genus. To investigate the role of *bvgS* in nutrient-induced filamentation, we conducted transcriptomics and found that *bvgS* mutation resulted in loss of responsiveness to highly-enriched media, especially in genes related to nutrient uptake and metabolism. The most dysregulated gene in the *bvgS* mutant encoded for succinyl-CoA:acetate CoA-transferase (SCACT) and we were able to regulate filamentation with exogenous metabolites up and downstream of this enzyme. These data suggest that *bvgS* regulates nutrient-induced filamentation by controlling metabolic capacity. Overall, we found that the virulence regulator *bvgS* is required for nutrient-induced filamentation in *B. avium*, suggesting there may be conservation in Bordetellae for utilizing this morphological change as a virulence phenotype.

## Introduction

The Bordetella genus comprises small Gram-negative bacteria, with the most notable being *Bordetella pertussis*, the causative agent of whooping cough in humans (1). The genus displays significant evolutionary plasticity, encompassing species that cause infection in animals or thrive in various environmental niches (2, 3). Through environmental sampling of wild nematodes, we previously identified a new species, *Bordetella atropi*, infecting the intestinal epithelia of the microscopic nematode *Oscheius tipulae. B. atropi* utilizes filamentation as a nutrient-induced virulence phenotype, whereby the bacteria detect host cell invasion to change morphology from coccobacilli to long filaments. These bacterial filaments are capable of invading neighboring intestinal epithelial cells in a novel intracellular cell-to-cell spreading mechanism (4). Filamentation by *B. atropi* is regulated by the glucolipid pathway in enriched nutrient environments, whereby excess UDP-glucose and the enzyme OpgH co-localize to constitutively inhibit FtsZ ring formation, leading to prolonged inhibition of binary division and filamentation (4, 5). Our prior results suggest that nutrient-sensing pathways, possibly influenced by host metabolites, can play a role in bacterial filamentation, and we wanted to investigate the conservation of nutrient-induced filamentation in Bordetellae.

Environmental signals and metabolite sensing significantly impact the virulence outcome in other *Bordetella* species. Despite evolutionary divergence, virulence in Bordetellae is predominantly governed by the highly conserved BvgAS master virulence regulator (6). This two-component system made up of BvgS (a sensor kinase) and BvgA (a response regulator) enables the bacteria to establish virulence within their hosts which allows the bacteria to switch between a virulent Bvg(+) and an avirulent/environmental Bvg(−) mode (7). Low temperatures, magnesium sulfate, and nicotinic acid were found to induce *B. pertussis* and *Bordetella bronchiseptica* into a Bvg(−) mode. By contrast, high temperatures or exposure to a host environment were found to induce a Bvg(+) mode (8). Both Bvg(−) and Bvg(+) modes are marked by changes in intracellular levels of phosphorylated BvgA, where high levels of BvgA∼P activate the transcription of virulence-associated genes (vags) (9), and low levels promote the transcription of virulence-repressed genes (vrgs) (7).

Microbial filamentation as a virulence mechanism has historically been associated with fungal hyphae, as seen in *Candida albicans*, which utilizes filamentation to disseminate throughout host tissues (10). In Gram-negative bacteria, filamentation is predominantly studied as a stress response, characterized by cell elongation without subsequent septation or division (11). Stress-induced filamentation is an adaptive strategy for survival and is known to be triggered by certain environmental contexts such as nutrient deprivation, UV-damage, or DNA-damaging antibiotics, as seen in *Bacillus subtilis* and *Escherichia coli* (12, 13). However, bacterial filamentation has also been proposed to be a potential virulence phenotype, with intracellular filamentation seen in clinically-relevant bacterial pathogens. For example, uropathogenic *E. coli* (UPEC) filaments inside umbrella cells of the urogenital tract, *Yersinia pestis* filaments in macrophages in vitro, and an isolate of *Salmonella typhimurium* was observed to filament in a culture of human melanocyte (14–16). The role these intracellular filaments play in virulence is largely unknown but is thought to be related to oxidative stress and resistance to phagocytes (17, 18). Given the discovery of *B. atropi*, it is possible that some intracellular bacteria may utilize nutrient sensing for induction of filamentation in vivo.

Within the Bordetella genus, *Bordetella avium* was previously found to filament in response to a change in nutrients in vitro, with the bacteria displaying a filamentous phenotype when shifted from standard growth media to enriched media (19). *B. avium* causes respiratory infections in birds (Bordetellosis) and contributes to economic losses in agriculture (20). We conducted a nutrient-based filamentation screen in *B. avium* and identified *bvgS* to be involved in regulating filamentation in enriched media. We conducted transcriptome comparisons between filamentous and non-filamentous *B. avium* and found that *bvgS* regulates several metabolic pathways, particularly those with TCA cycle intermediates, previously not linked to virulence in Bordetellae. We focused on the enzyme succinyl-CoA:acetate CoA-transferase (SCACT), which was the most regulated gene by *bvgS* in enriched media and found that exogenous metabolites upstream and downstream of this enzyme could regulate filamentation. Our results suggest that BvgS potentially allows for a metabolic shift, allocating resources for filamentation in nutrient-rich environments, and for survival in nutrient-depleted environments. Overall, our results provide valuable insights into the connection between bacterial metabolism and virulence, shedding light on genes not normally associated with a Bvg(+) or Bvg(−) mode in *Bordetella* species.

## Results

### *Bordetella avium* filaments in highly-enriched media

We had previously discovered that the bacterial species *B. atropi* uses nutrient-induced filamentation as a novel virulence phenotype, using filaments for intracellular spreading in the nematode *O. tipulae*. Prior research on the related bacterium *B. avium* found that this species could filament when switched from a standard rich media (tryptic soy broth, TSB) to a highly enriched media (terrific broth, TB) (19). However, this work did not determine the cause of the filamentous phenotype or any relationship between the phenotype and virulence. We validated this work and established a consistent induction protocol for generating filamentous *B. avium* using highly-enriched media. First, we looked at *B. avium* morphology in a chemically defined media (CDM), which would have the least nutrient density of the media we tested, and found the bacteria displays a majority coccobacilli morphology (**Fig. 1A**) (21). Then, we found that *B. avium* grown in TSB displays a mixed coccobacilli and short filament phenotype with occasional filaments, as seen previously (**Fig. 1B**). When switched from TSB to TB, the majority of *B. avium* bacteria display a dramatical increase in length, appearing as long filaments (**Fig. 1C**). However, a small proportion of bacteria in TB were found to be non-filamentous coccobacilli, suggesting that there is stochasticity in the bacterial response to increased nutrient availability.

**Figure 1.**
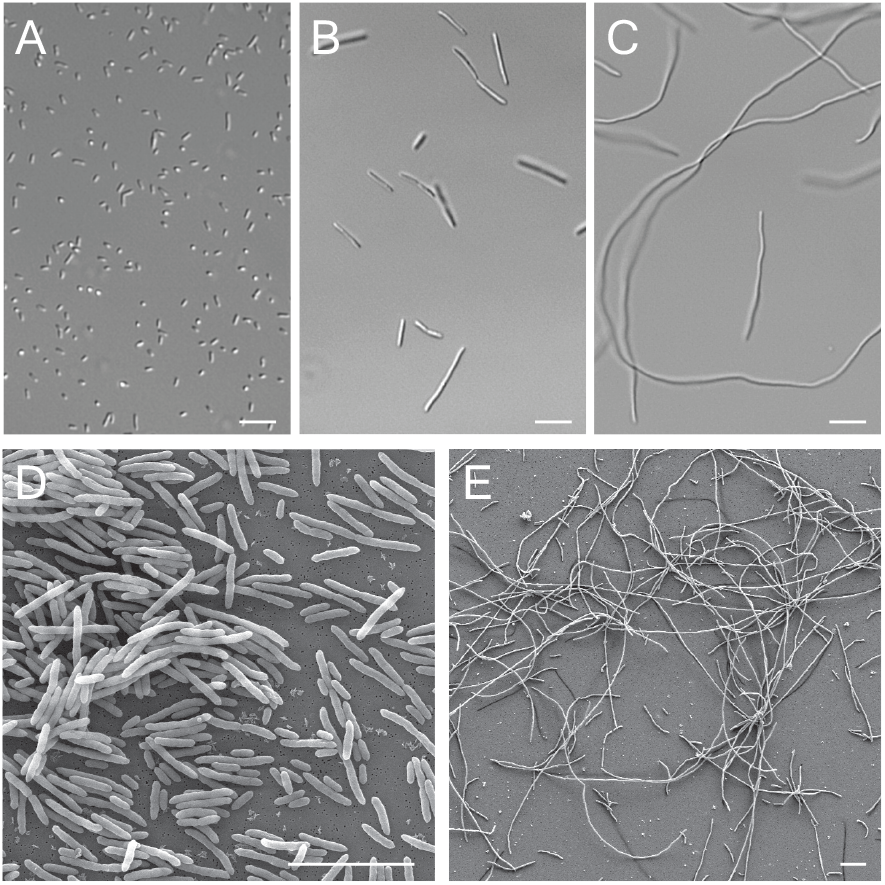
Filamentation of *Bordetella avium* in highly-enriched media. (**A**) Light microscopy image of *B. avium* grown in chemically defined media (CDM (**B**) Light microscopy image of *B. avium* grown in tryptic soy broth (TSB). (**C**) Light microscopy image of *B. avium* grown in terrific broth (TB). (**D**) Scanning electron microscopy (SEM) image of *B. avium* grown in TSB. (**E**) SEM image of *B. avium* grown in TB. Scale bars = 10 µm.

We also used scanning electron microscopy (SEM) to look at *B. avium* grown in TSB or TB. We observed mostly non-filamentous coccobacilli in TSB and long filaments grea ter than 50 mm in length in TB. The surface of both morphotypes of *B. avium* appear rough rather than smooth, similar to what is seen in TEM images of *B. atropi* in vivo (4) (**Fig. 1D, E**). We observed that septation of filamentous *B. avium* mostly occurs at the midline of each filament, generating a new midline for each section following the initiation of the first (**Supp. Fig. 1A**). Strikingly, we observed that some of the *B. avium* filaments showed internal branching, a phenomenon most associated with Gram-po sitive bacteria, like Nocardia and Actinomyces (22) (**Sup. Fig. 1B-C**). Overall, we validated that *B. avium* was capable of undergoing nutrient-induced filamentation, similar to *B. atropi*, and found a method to consistently induce the bacteria to filament in vitro.

### Isolation of *B. avium* mutants that fail to filament when switched into highly-enriched media

To identify any genes that regulate filamentation in *B. avium*, we conducted a selection-based screen previously used to isolate non-filamentous mutants of *B. atropi*. As described above, *B. avium* will consistently filament when grown in T B after being transferred from TSB. We utilized this paradigm to induce filamentation then filter out filamentous bacteria with a 5 µm filter before the flowthrough was reinoculated into TSB. This procedure was conducted with two parallel cultures and repeated 9-10 times to yield two independent non-filamentous mutant strains (**Fig. 2A**). *B. avium* mutant 10a-1 and mutant 9b-6 from these parallel screens were validated to not filament when grown in TB, with an average bacterial length of 2 and 2.5 µm in TB (**Fig. 2B, C**). The bacterial length of the mutants were significa ntly smaller than WT *B. avium* grown in TB (mean=40 µm), as well as significantly smaller than WT *B. avium* grown in TSB (mean=11 µm) (**Fig. 2D**). To test whether the lack of filamentation might be due to impaired growth kinetics, we compared the growth rate of mutant 9b-6 to WT *B. avium* over the course of 48 hours in TSB, TB, and CDM. We decided to focus on the growth kinetics of mutant 9b-6, as mutant 10 a-1 presented other phenotypic traits such as agglutination that made it harder to measure (**see Fig. 2C**). We found that both the mutant 9b-6 and WT *B. avium* have identical growth kinetics in CDM. Similarly, mutant 9b-6 and WT *B. avium* have identical growth kinetic in TSB and TB from 0-15 hours. However, after 15 hours WT *B. avium* has a decreased OD_600_ compared to mutant 9b-6 (**Fig. 2E**), likely due to sedimentation of the large bacterial filaments and/or inconsistent absorption by filaments.

**Figure 2.**
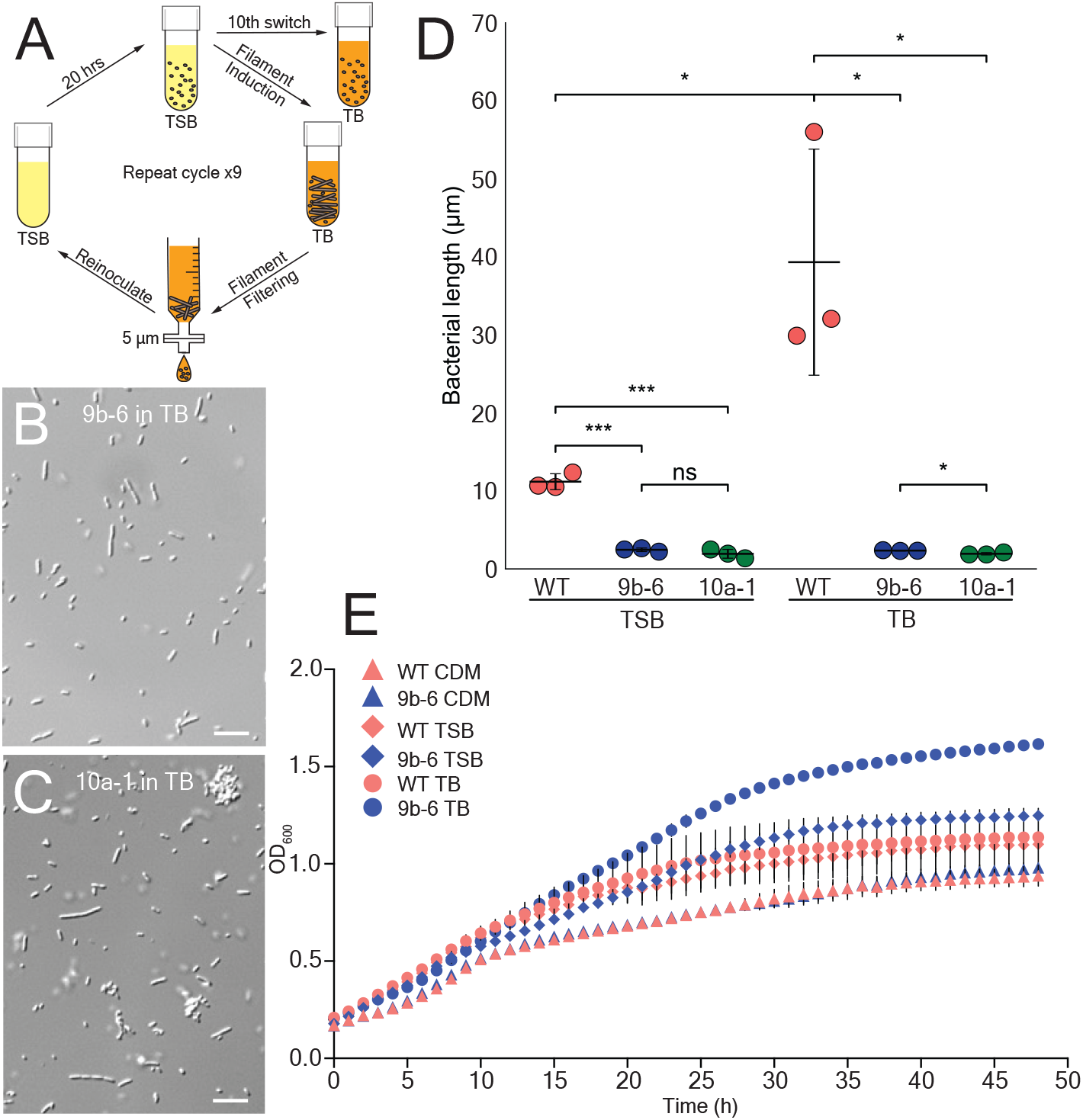
Isolation of non-filamentous mutants of *B. avium*. (**A**) Schematic illustrating filamentation induction of *B. avium* through switching from TSB to TB, followed by a filtration-based mutant isolation process. Non-filamentous bacteria were reintroduced into TSB, with successive filtration until the filamentous population in TB was reduced to less than 5%. (**B**) Light microscopy image of mutant 9b-6 grown in TB after switching. (**C**) Light microscopy image of mutant 10a-1 grown in TB after switching. Scale bars = 10 µm. (**D**) Scatter plot comparing the average bacterial length per replicate of WT *B. avium* (red) and non-filamentous mutants 9b-6 (blue) and 10a-1 (green) in TSB and TB. Experiment was done in triplicate (N=3), n=200 bacteria per replicate, p<0.05 (*), p< 0.001 (***), ns=not significant, by unpaired two-tailed t-test. Error bars represent SD. (**E**) OD600 growth curves of WT *B. avium* and non-filamentous mutant 9b-6 in chemically defined medium (CDM), TSB, and TB. N=3, error bars represent SD.

### Sensor kinase (BvgS) of the conserved master virulence regulator (BvgAS) regulates filamentation in vitro

To identify the gene(s) required for filamentation, we conducted long-range, whole genome sequencing of mutants 10a-1 and 9b-6, and the WT strain. Mutant 9b-6 had a total of two polymorphisms in predicted protein coding genes which would change the amino acid sequence, including a 30-nucleotide insertion. Mutant 10a-1 had a total of five non-synonymous polymorphisms in predicted protein coding genes. Interestingly, both mutants had different allelic mutations in the same gene, *bvgS*, of the Bordetella conserved master virulence regulator, BvgAS. Mutant 10a-1 had a non-synonymous point mutation *bvgS*^D580Y^, and mutant 9b-6 had a ten amino acid insertion, *bvgS*: p.R687_V688insLLPYSDSLGR (**Fig. 3A, B**). These mutations in *bvgS* are in or around the Per-Arnt-Sim (PAS) domain and the PAS-associated C-terminal domain (PAC) motif. The PAS domain is associated with sensing environmental changes and signal transduction (**Fig. 3C**) (23). Both the PAS domain and the PAC motif are found in a wide range of signaling transmembrane proteins and they typically lie in the cytoplasmic portion (24). The PAS domain and PAC motif were previously identified to be in the cytoplasmic portion of BvgS in *B. bronchiseptica*, and mutations of these domains were linked to a constitutive avirulent phase (25).

**Figure 3.**
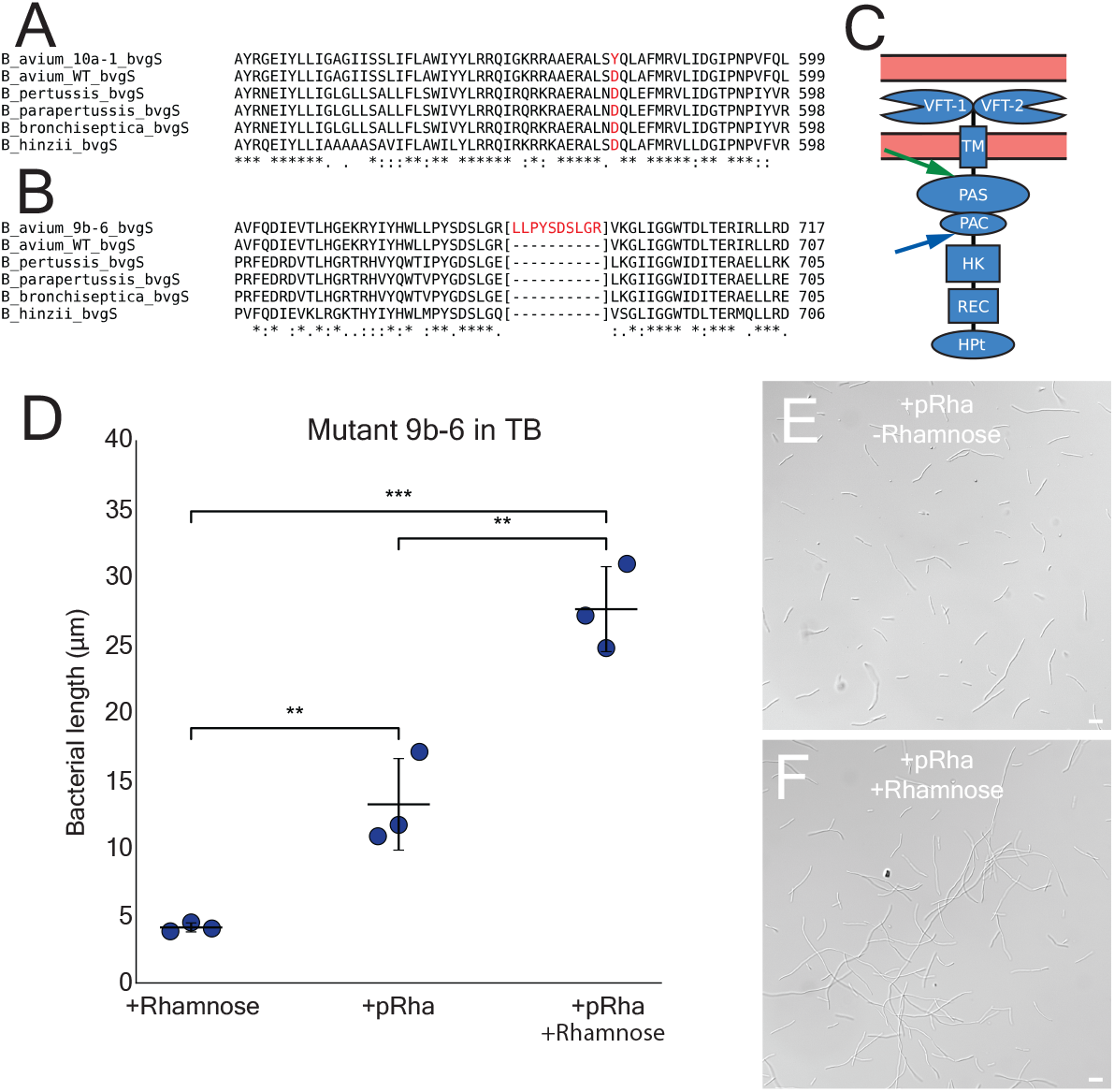
Identification and complementation of *Bordetella avium* non-filamentous mutants. (**A-B**) CLUSTAL O multiple sequence alignment of the Bordetella virulence regulator protein BvgS, showing mutations in non-filamentous mutants. Mutant 10a-1 (A) contains a point mutation (D580Y), and mutant 9b-6 (B) presents a repeat ten-amino acid insertion (p.R687_V688insLLPYSDSLGR). (**C**) Predicted diagram of the protein BvgS with known domains, from top to bottom, Venus Flytrap 1 & 2, transmembrane, Per-Arnt-Sim, PAS-associated C-terminal motif, histidine kinase, receiver, histidine phosphotransfer. Figure based on similar structures presented in *B. bronchiseptica* (26), with a green arrow indicating the location of the point mutation (D580Y) found in mutant 10a-1, and blue arrow indicating the location of the insertion (p.R687_V688insLLPYSDSLGR) found in mutant 9b-6. (**D**) Scatter plot comparing the average bacterial length per replicate of non-filamentous mutant 9b-6 complemented with WT *bvgS*, with and without induction of plasmid using rhamnose (200 µg/ml). Experiment was done in triplicate (N=3), n=200 bacteria/replicate, p<0.01 (**), p<0.001 (***) by unpaired two-tailed t-test. Error bars represent SD. (**E**) Representative light microscopy image of mutant 9b-6 with pANT-4-*bvgS* grown in TB without rhamnose. (**F**) Representative light microscopy image of mutant 9b-6 with pANT-4-*bvgS* grown in TB with rhamnose. Scale bars = 10 µm.

In order to validate that *bvgS* is required for filamentation on highly-enriched media, we complemented mutant 9b-6 with a WT copy of *bvgS* on a rhamnose inducible plasmid. Untransformed 9b-6 grown in rhamnose showed an average bacterial length of ∼5 mm. *BvgS* complemented 9b-6 grown in rhamnose displayed a heterogenous population of bacterial morphologies including coccobacilli and short filaments, with an average bacterial length of 13 µm. This increase in filamentation without the addition of rhamnose is likely due to leaky expression due to endogenous rhamnose biosynthesis by *B. avium* (**Fig. 3D, E**). Finally, *bvgS* complement 9b-6 with rhamnose displayed a majority of large filaments, with an average bacterial length of 28 µm when grown in TB after switching (**Fig. 3D, F**). Altogether, these results show that the loss of nutrient-induced filamentation seen in mutant 9b-6 can be rescued with a WT copy of *bvgS*.

### Transcriptomics analysis of WT and mutant 9b-6 strains in TB and CDM

In order to identify the genes that are controlled by *bvgS* when switched into highly-enriched media, we conducted transcriptomics on WT *B. avium* and non-filamentous mutant 9b-6 grown in a non-filamenting (CDM) and filamenting conditions (TB). The *B. avium* genome contains 3,483 protein coding genes. For WT *B. avium*, we found that the expression of 473 genes were significantly upregulated by two-fold or more in TB compared to CDM with a false discovery rate (FDR) of 5%. Similarly, 447 genes were significantly downregulated by two-fold or more **(Fig. 4A)**. By contrast, mutant 9b-6 upregulated 362 genes after switching into TB and downregulated 324 genes **(Fig. 4B)**.

**Figure 4.**
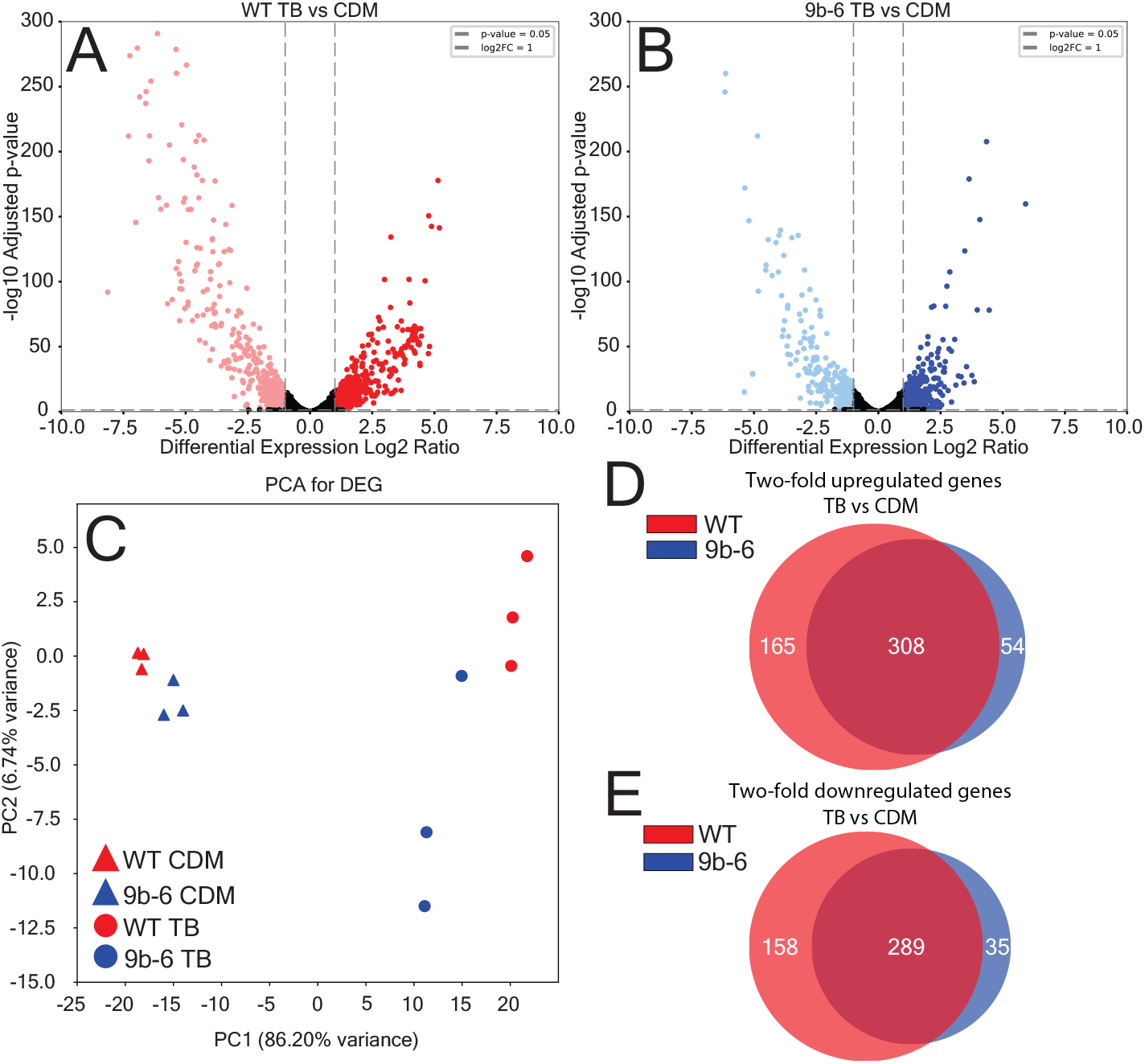
Comparative transcriptomic analysis of WT and mutant 9b-6 strains in TB and CDM. (**A-B**) Volcano plots showing the distribution of DEGs between WT (A, left) and mutant 9b-6 (B, right) strains in TB after switching from CDM. Each plot represents a gene, plotted by its -log10 adjusted p-value and Log2 fold change (FC). DEGs with a Log2FC ≥ 1 or ≤ -1 are highlighted in red and pale red for WT, and blue and pale blue for mutant 9b-6. The dashed lines denote the statistical significance threshold of p = 0.05 and Log2FC = ±1. DEGs outside of the threshold are presented as black. (**C**) Principal component analysis (PCA) of DEGs for WT and 9b-6 strains in both TB and CDM conditions. The separation of the samples along the principal components (PC1 and PC2) demonstrates the transcriptomic differences between the conditions and replicates, respectively. (**D-E**) Venn diagrams of upregulated (D, Log2FC > 1) and downregulated (E, Log2FC < -1) DEGs in the WT (red) and mutant 9b-6 (blue) strains in TB after switching. Shared DEGs are shown in the overlap, while strain-specific DEGs are in non-overlapping regions.

We performed a principal component analysis (PCA) on the differentially expressed genes (DEGs) to assess the variance between WT and mutant 9b-6 strains under the two different growth conditions, CDM and TB. PCA was conducted on all the samples, with the replicates clustering near each other **(Fig. 4C)**. Over 90% of the variance in the samples were represented in the 2-dimensional plot, with 86.20% of the variance along the first principal component (PC1) and 6.74% along the second principal component (PC2). There is a clear separation between the TB and CDM conditions along PC1, indicating that the growth environment has a dominant influence on variance in gene expression. Additionally, within each condition, there is a distinct separation between WT and mutant 9b-6 along PC1. Interestingly, the WT strain shows a larger separation between CDM and TB along PC1 compared to mutant 9b-6, indicating a greater response by WT bacteria upon switching to highly-enriched media.

We further analyzed the DEGs by comparing the up- and downregulated genes between the WT and mutant 9b-6 strains in TB after switching to identify genes that were either commonly or uniquely expressed in each strain. We compared significant DEGs from the WT and mutant 9b-6 strains that were upregulated greater than two-fold. Out of these, 165 genes were uniquely upregulated in the WT strain (shown in red), and 54 genes were uniquely upregulated in the mutant 9b-6 strain (shown in blue) (**Fig. 4D**). Similarly, we compared significant DEGs from the WT and mutant 9b-6 strains that were downregulated greater than two-fold. Out of these genes, 158 genes were uniquely downregulated in the WT strain and 35 genes were uniquely downregulated in the mutant 9b-6 strain **(Fig. 4E)**. These results highlight that a large set of genes are regulated by both strains, but a greater number of uniquely expressed genes were found in the WT strain compared to the mutant 9b-6 strain. This data indicates that WT bacteria is more responsive to a highly-enriched media, implicating *bvgS* in regulating transcription after nutrient changes.

### Functional characterization of gene expression changes and implications of metabolism on filamentation

To investigate the functional implications of differential gene expression in the mutant 9b-6 strain, we performed a Gene Ontology (GO) Slim analysis, focusing on br oad categories of enriched molecular functions and biological processes. The analysis was conducted on unique WT or mutant 9b-6 DEGs with a greater than 2-fold increase in gene expression. Overall, we observed significant enrichment in the WT strain in GO-term categories associated with transferase activity, hydrolase activity, transport activity, transmembrane transport, and signaling.

These categories were largely absent in mutant 9b-6, suggesting a dysregulation of these processes when switched to TB (**Fig. 5A-B**). These findings suggest that the WT strain exhibits greater adaptability in regulating metabolic pathways in response to an increase in nutrients compared to mutant 9b-6.

**Figure 5.**
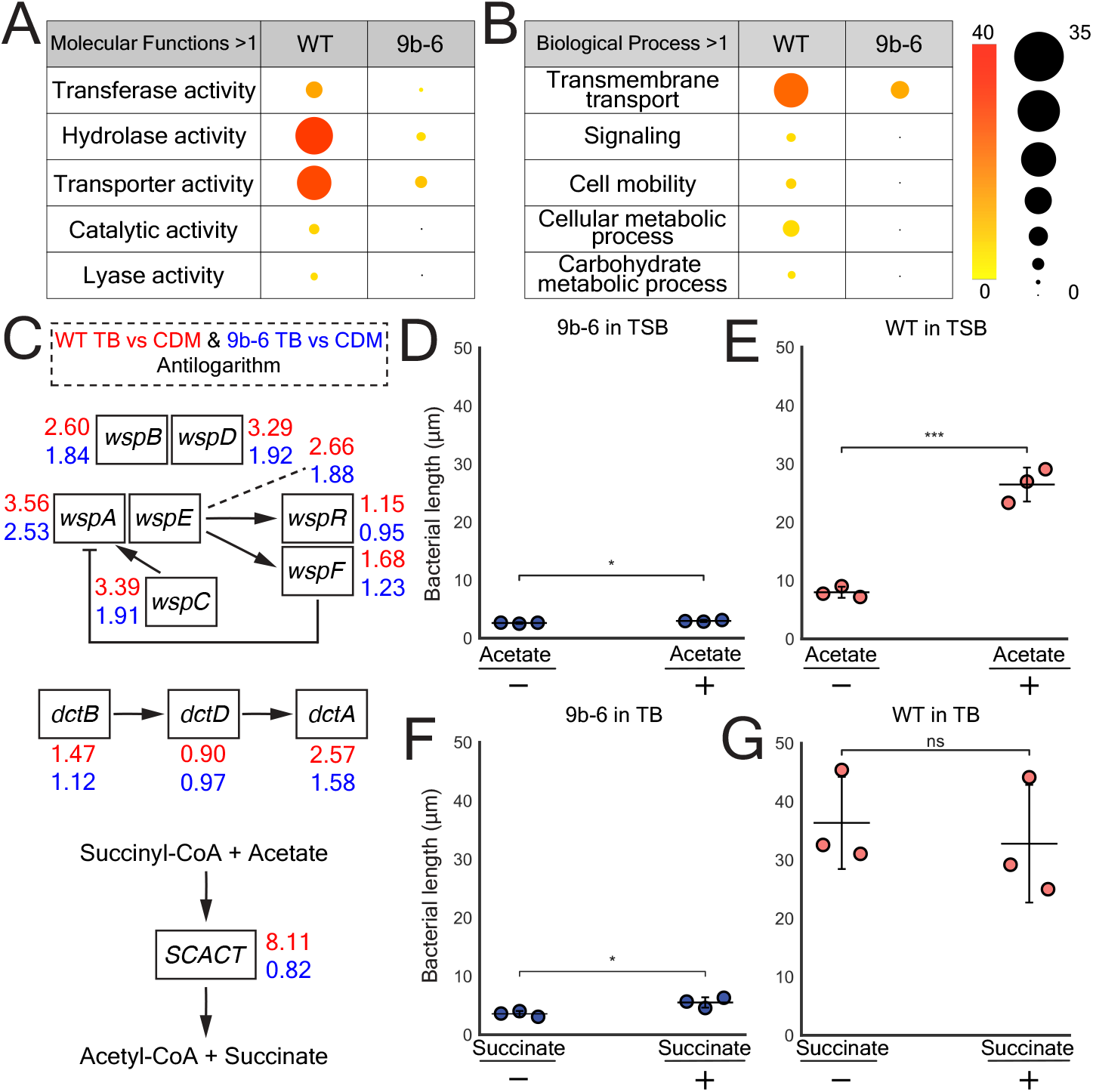
GO-term enrichment DEGs in WT and mutant 9b-6 *Bordetella avium* when grown in TB and filamentation rescue using metabolites. (**A-B**) Bubble tables showing significantly enriched GO-terms for DEGs with Log2FC ≥ 1 in WT and mutant *B. avium* grown in TB after switching. GO-term enrichment analysis reveals significant upregulation of genes associated with enzymatic activity and transport activity, in WT *B. avium* in comparison to mutant 9b-6. The color of the bubbles represents the node-score (significance), and the size of the bubbles corresponds to the number of sequences associated with each GO-term. (**C**) Diagram of three separate pathways singled out based on enriched GO-terms and analysis through KEGG pathway analysis showing fold-change in TB vs CDM gene expression. WT values are shown in red and mutant 9b-6 in blue. Wsp operon involved in regulating biofilm formation and surface associated behaviors (*top*). Dct operon involved in transport and regulation of C4-dicarboxylates (*middle*). The enzyme SCACT allowing for succinyl-CoA to be directly converted to acetyl-CoA and succinate via acetate (*bottom*). (**D-E**) Scatter plot comparing the average bacterial length per replicate of non-filamentous mutant 9b-6 (D) and WT *B. avium* (E) grown in TSB supplemented with 20mM sodium acetate. Experiment was done in triplicate (N=3), n=200/replicate, p<0.05 (*), p<0.001 (***) by unpaired two-tailed t-test. Error bars represent SD. (**F-G**) Scatter plot comparing the average bacterial length per replicate of non-filamentous mutant 9b-6 (F) and WT *B. avium* (G) grown in TB supplemented with 1mM succinate. Experiment was done in triplicate (N=3), n=200/replicate, p<0.05 (*), ns=not significant, by unpaired two-tailed t-test. Error bars represent SD.

To further understand the gene expression changes in WT compared to mutant 9b-6 strains, we conducted a KEGG pathway analysis on all genes with two-fold or greater changes after switching to TB. Several pathways demonstrated increased activity across both strains, most notably the uptake and processing of branched chain amino acids through the Liv operon. However, WT bacteria specifically upregulated certain pathways compared to mutant 9b-6 after switching to TB. Specifically, the Wsp operon, encoding a chemotaxis like signaling pathway involved in surface associated behavior and regulation of biofilm formation, was comprehensively upregulated (**Fig. 5C**), suggesting that *B. avium* filamentation could play a role biofilm formation in nutrient-rich niches. Further, the WT strain also exhibited notable upregulation of the C4-dicarboxylate transporter (*dctA*) gene, encoding a membrane transport protein, suggesting that a mutation in the senor kinase BvgS affects the uptake of 4-carbon metabolites such as succinate and fumarate in highly-enriched media (**see Fig. 5C**). Finally, the gene encoding succinyl-CoA:acetate-CoA transferase (SCACT) exhibited a greater than 8-fold increase in the WT strain compared to no significant difference in mutant 9b-6 grown in TB, representing the largest difference in gene expression between these two strains (**see Fig. 5C**). SCACT converts acetate and succinyl-CoA into succinate and acetyl-CoA.

Overall, our KEGG analysis indicated that *bvgS* may play a role in detecting and/or processing metabolites like succinate and acetate in *B. avium*, leading to bacterial filamentation. To test this, we first focused on the acetate, a metabolite upstream of SCACT. We found that supplementing WT *B. avium* in TSB with sodium acetate (20 mM) resulted in a significant increase in filamentation, from a mean of 8.0 µm to 26.5 µm, while mutant 9b-6 showed a modest increase from 2.6 µm to 2.9 µm (**Fig. 5D-E**). Given the increase in SCACT gene expression in WT bacteria in TB, this data suggests that acetate processing is important for *bvgS-*dependent induction of filamentation. Next, we focused on succinate, a metabolite downstream of SCACT and in the TCA cycle. When succinate (1 mM) was supplemented to the mutant 9b-6 and WT *B. avium* s trains grown in TB, we saw that succinate could partially rescue filamentation in mutant 9b-6 in enriched media, increasing from 3.6 µm to 5.5 µm, while the WT strain showed no change, most likely due to an already saturated level of succinate (**Fig. 5F-G**). This data suggests that succinate levels in *B. avium* may play a role in induction of filamentation. We further tested if direct downstream inhibition of succinate dehydrogenase which facilitates succinate being catalyzed into fumarate in the TCA cycle could block filamentation. Supplementing mutant 9b-6 grown in TB + malonate (80 mM) resulted in no significant changes (**Supp. Fig. 2A**). Strikingly, supplementing WT *B. avium* grown in TB + malonate (80 mM) resulted in near complete loss of filamentation, suggesting that inhibition of succinate dehydrogenase disrupts filamentation (**Supp. Fig. 2B**). We further tested if manipulation of the TCA cycle directly downstream of succinate could affect filamentation. Supplementing mutant 9b-6 grown in TB with fumarate or malate largely resulted in no significant changes in filamentation, except 100 mM fumarate resulted in mild de crease in bacterial length (**Supp. Fig. 2C**). In contrast, supplementing WT *B. avium* grown in TB with fumarate or malate largely resulted in a significant reduction of filamentation (**Supp. Fig. 2C**). As such, other TCA cycle metabolites (besides succinate) do not play a role inducing *B. avium* filamentation in enriched media. Instead, manipulation of the TCA cycle with fumarate, malate, and malonate results in filamentation inhibition, perhaps due to indirect inhibition of enzymatic activity following upstream metabolic saturation. Taken together, these findings indicate that the WT strain exhibits greater adaptability in nutrient rich environments, with *bvgS* control of SCACT gene expression likely playing a partial role in filamentation induction in nutrient rich conditions.

## Discussion

Our study demonstrates that *B. avium* undergoes filamentation in the nutrient-rich conditions associated with TB, suggesting a link between nutrient availability and bacterial morphology. While bacterial filamentation is often associated with stress responses, our findings align with reports from the related species *B. atropi*, where filamentation was required for a virulence phenotype, cell-to-cell spreading in the nematode *O. tipulae*. In *B. avium*, we established a reproducible protocol for inducing filamentation and showed that filamentous phenotypes are predominant and abundant in TB. Scanning electron microscopy revealed unique features, like occasional branching (see Supplemental Fig. 1A), further underscoring the morphological adaptations of *B. avium* in response to nutrient conditions. We further identified that mutations in the sensor kinase gene *bvgS*, part of the BvgAS master virulence regulatory system, was required for filamentation in nutrient rich conditions and regulated a set of gene likely associated with nutrient uptake and processing. We validated that SCACT likely plays a part in metabolic regulation, ultimately leading to filamentation in nutrient-rich conditions like TB. These discoveries implicate BvgS as a key regulator of filamentation in *B. avium* and implicate this phenotype in playing a role in virulence.

Gram-negative bacteria like *Escherichia coli, Salmonella enterica*, and *Yersinia pestis* have previously been identified to utilize filamentation as a potential virulence phenotype (14–16). Additionally, the observation of nutrient-induced intracellular filamentation of *B. atropi* plays an in vivo role in infection supports the idea that filamentation can be a pathogenic phenotype with a clear role in virulence. Our data suggest that nutrient-induced filamentation in *B. avium* is controlled by the two-component master virulence regulator, BvgAS, enabling the bacteria to sense nutrient shifts that signify either a rich or poor environment (27). It has been known that environmental cues allow Bordetella species to adapt to different niches, with the bacteria switching between a pathogenic Bvg(+) mode when in a host and a survival-oriented Bvg(−) mode when in the environment (7). However, the triggers for this adaptive behavior are not fully understood. Our data suggest a dynamic relationship between nutrient availability and the bacterial decision to induce virulence through BvgAS, such that *B. avium* may differentiate between a nutrient-rich niche in the host from a nutrient-poor niche. As such, an abundant nutrient source would allow for accumulation of internal metabolites to support and initiate a virulence phenotype (filamentation), making the potential trigger an internal non-host derived stimuli rather than a host derived external stimuli. Similar behaviors have been identified in quorum sensing of *Vibrio cholerae*, where autoinducers regulate virulence and biofilm formation (28). While the relationship between BvgS and filamentation of *B. avium* suggests a Bvg(+) mode, previous work on *B. bronchiseptica* show that mutations in the PAS domain and accompanied PAC motif yields a locked Bvg(−) mutant (25). These differences may be due to species-specific effects or the nature or the alleles in these domains, as we were unable to determine whether the non-filamentous mutant 9b-6 is locked in Bvg(−).

Currently, the in vivo function of nutrient-induced filamentation in *B. avium* remains unknown. It is possible that a nutrient-rich environment in a host may allow *B. avium* to allocate resources toward filamentation to support downstream processes such as proliferation, biofilm formation, and/or niche establishment (29). *BvgS-*dependent biofilm formation is supported by our finding of the upregulation of the Wsp operon under nutrient rich conditions. Interestingly, biofilm formation is linked to filamentation in Gram-negative bacteria like *E. coli* and *Xylella fastidiosa*, and is a well-known virulence phenotype (30, 31). Furthermore, our finding of the C4-dicarboylate protein DctA is upregulated in the WT strain and that the addition of succinate promotes bacterial length in the mutant 9b-6, suggests that succinate, or a steady and bountiful production of succinate via SCACT is crucial for filamentation. Succinate is a preferred carbon source for Bordetella and a common metabolite in host environments (32, 33). Additionally, succinate is a critical carbon source that can trigger biofilm formation by *Pseudomonas aeruginosa* in a cystic fibrosis lung and act as an intracellular trigger for virulence in *Salmonella* Typhimurium (34, 35)

We found that SCACT was the most dysregulated gene in the *bvgS* mutant, and we were able to regulate filamentation with exogenous acetate and succinate. The Bordetella genus presents immense plasticity, exemplified by the variety of species found in diverse niches, including animals, like humans, dogs, birds, and nematodes, as well as environmental niches, like contaminated soil and dechlorinating bioreactors (3). Many mammalian-infecting species utilize *bvgAS* to switch between a potentially harsh environmental niche and a host. Interestingly, SCACT is a key enzyme for adapting to an acetate-rich environment, acting as primary shunt for succinate production (36), and hosts generally produce an abundance of acetate as it is a common byproduct of fermentation and beta-oxidation (37). Therefore, it is possible that in the process of establishing infection, host acetate is utilized as a carbon source by *B. avium* to facilitate filamentation and potentially downstream biofilm formation. In fact, acetate, as well as TCA cycle intermediates and amino acids like glutamate and proline, are all known preferred carbon sources of *B. bronchiseptica* and *B. pertussis*, due to an inability to utilize carbohydrates and sugar alcohols as sole carbon sources (32, 33). There is also potential for *B. avium* co-infection of the lung, as avian pathogenic *E. coli* (APEC) triggers production of microbiota-derived acetate as a defense mechanism and protectant against persistence and development of APEC induced lung inflammation (38). Poultry having success in in mitigating persistence of infection of APEC through the use of acetate, provides a unique opportunity for *B. avium* to take advantage of excess acetate to develop a pathogenic phenotype.

In summary, we found that the master virulence regulatory system, BvgAS, controls nutrient-induced filamentation in *B. avium*, suggesting this virulence phenotype may be conserved beyond the nematode-infecting *B. atropi*. We identified SCACT, which allows for a metabolic shift in the TCA cycle, was regulated by BvgS, allowing *B. avium* to efficiently allocate resources towards either survival, or virulence through filamentation. While primarily associated with avian hosts, *B. avium* also has potential zoonotic implications, underscoring the importance of understanding the genetic control of genes involved in nutrient-induced filamentation (39). Insights gained in *B. avium* could broaden our overall understanding of filamentation as a virulence phenotype and aid in developing strategies against more clinically significant species such as *B. pertussis, B. parapertussis*, and *B. bronchiseptica* (40).

## Materials and Methods

### Bacterial strains and media

*B. avium* IPDH 591-77 (35086) was acquired from ATCC and maintained on TSB agar plates. *Escherichia coli* SM10 was acquired from ATCC and maintained on TSB and LB agar plates. CDM contains: Base solution: KH_2_PO_4_ 0.5 g, Sodium glutamate 10.72 g, NaCl 2.5 g, KCl 0.2 g, MgCl_2_ * 6H_2_O 0.1 g, Tris 1.525 g, α-Ketoglutaric acid 2.0 g, Sodium pyruvate 2.0 g, CaCl_2_ 20.0 mg, H_2_O to 909.0 mL, pH adjusted to 7.8 using NaOH. CDM supplement: Ascorbic acid 0.2 g, Glutathione 1.0 g, H_2_O to 100.0 mL. L-Cystine supplement: H2O 3.0 mL, HCl (11.6 M) 1.0 mL, L-Cystine 0.4 g, H_2_O to 100.0 mL. L-Proline supplement: L-Proline 3.0 g, H2O to 200.0 mL. L-Phenylalanine supplement: L-Phenylalanine 4.0 g, H_2_O to 400.0 mL. Vitamins: p-Aminobenzoic acid 10.0 mg, Biotin 1.0 mg, Folic acid 10.0 mg, Pyridoxyl HCl 80.0 mg, Riboflavin 40.0 mg, Thiamine HCl 40.0 mg, Calcium pantothenate 80.0 mg, Nicotinamide 200.0 mg, H_2_O 800.0 mL, NaOH (2.5 M) until solution clears, H_2_O to a final volume of 1000.0 mL. Final media preparation: Base solution 909.0 mL, L-Cystine stock 10.0 mL, L-Proline stock 16.0 mL, CDM supplement 10.0 mL, L-Phenylalanine stock 40.0 mL, Vitamins 10.0 mL. Final sterilization through ultrafiltration (0.2 μm) (21). TSB contains: Tryptone 15.0 g/L, Soybean peptone 5.0 g/L, NaCl 5.0 g/L, K2HPO4 2.5 g/L, Glucose 2.5 g/L. TB contains: Tryptone 20.0 g/L, Yeast extract 24.0 g/L, Glycerol 4.0 mL/L, KH2PO4 2.31 g/L, K2HPO4 12.54 g/L.

### Filamentation induction

All *B. avium* strains were grown at 37°C overnight while shaking at 200 RPM in either CDM, TSB, or TB, unless specified otherwise. Nutrient-induced filamentation of *B. avium* was achieved by transferring 10 μL of bacteria grown in CDM or TSB into 5 mL TB and incubated at 37°C overnight while shaking at 200 RPM. When supplementing with metabolites, 0.5 M stock solutions of succinic acid (succinate), sodium malonate, or sodium acetate were dissolved in water and filter sterilized using 0.2 μm PES filters. Metabolites were used to supplement TB or TSB at 1 mM, 80 mM, and 20 m for succinate, malonate, and acetate respectively.

### Scanning electron microscopy

WT *B. avium* and *B. avium* 9b-6 were grown in TSB overnight and later transferred to TSB or TB. Bacteria were pelleted and fixed in a 1:1 mixture of 4 % glutaraldehyde and 0.1M cacodylate buffer for 1 hour. Then, samples were filtered through a 0.1 μm filter, directly followed by 0.1 M cacodylate buffer wash. The filter containing bacteria was removed from the syringe and placed facing up in a 6 well plate with cacodylate buffer and vapor fixed with OsO4 for 1 hour. Samples were dried using consecutive EtOH washes with 50%, 70%, 2X 95%, and 2X 100% EtOH at 15 minutes each. Samples were left overnight to dry in hexamethyldisilazane. Filters were mounted on carbon double sticky tape and coated in 6 nm platinum before visualization through SEM.

### Non-filamentous mutant screen

Two parallel colonies of WT *B. avium* were induced to filament as above. Filament containing samples in TB were filtered using a 5 μm filter. Bacterial flowthrough was reinoculated into TSB and grown overnight before another round of filamentation induction in TB. The bacteria were filtered a total of 9-10 times until <5% filamentation was visualized in the population. The final flow-through of the two replicates were plated onto TSB plates and grown overnight at 37°C. Colonies were picked and validated to not filament in TB after switching in vitro.

### Whole genome sequencing and analysis

WT, mutant 9b-6, and 10a-1 were cultured in TSB and DNA was purified using the Qiagen DNeasy Blood and Tissue Kit. Whole genome sequencing, assembly, and analysis was conducted as described previously using Oxford Nanopore R10.4.1 flow cells after v14 Library Preparation by Plasmidsaurus (41). Reads from mutant 9b-6 and 10a-1 were mapped onto the assembled and annotated WT genome using Geneious Prime. Mutations were identified by scanning across the genome using Geneious Prime.

### Cloning

The WT copy of *bvgS* was cloned into the pANT4_pRha_GFP plasmid using restriction enzyme cloning. The pANT4 plasmid with the rhamnose promoter was designed by Dr. Anca Segall and engineered by Dr. Kyle Malter using sequences of pANT4 and pRha-67 (42, 43). The promoter provides a better rheostat control of gene expression (44). This pANT4-based expression system has been used in Achromobacter xylosoxidans for complementation experiments, where expression is constitutive (Ryan Rowe, Hamza Hajama, and Anca Segall, ms. in preparation. Briefly, the entire *bvgS* gene was amplified from WT *B. avium* genomic DNA using primers (5’-TCGGTACCCGGGGATCC<u>TCTAGA</u>GTGGGCTTAAA GAGCTGCTGGAG-3’ and 5’ GCATGCAAGCTTGGCTGTTTTGGTCAGGCACGCGACGCCAAC 3’). Primers were designed with overhangs containing restriction enzyme digestion sites. The PCR product was extracted after gel electrophoresis using Zymoclean™ Gel DNA Recovery Kit and then cleaned using DNA Clean & ConcentratorTM5. Plasmid pANT4_pRha_GFP was maintained in SM10 and extracted using Zyppy™ Plasmid Miniprep Kit and cleaned using DNA Clean & ConcentratorTM-5. PCR product and plasmid were digested using XbaI and HindIII, with the digested plasmid purified from a 0.7% agarone gel. The PCR product and plasmid were ligated using T4 DNA Ligase according the manufacturer’s guidelines and transformed into chemically competent SM10. Plasmid pANT4_pRha*_bvgS* was verified through whole plasmid sequencing.

### Electroporation

*B. avium* mutant 9b-6 was grown in Super Optimal Broth (SOB) to an optical density (OD600) of 0.5-0.8. All material, 10% glycerol and bacterial cultures were chilled on ice for 30 minutes. Bacteria were pelleted at 4200 g at 4°C and pellet was washed in 10% glycerol. Centrifugation and wash step was repeated four more times. The final bacterial pellet was suspended in residual 10% glycerol. pANT4_pRha_bvgS was purified using Zyppy™ Plasmid Miniprep Kit, and Qiagen HISPEED® Plasmid Midi Kit. For electroporation of *E. coli, ∼*500 ng of pANT4_pRha_bvgS was loaded into 50 μL of competent cells and allowed to incubate on ice for two minutes, before transfer into a chilled 2 mm electroporation cuvette, and electroporated using BIORAD MicroPulser™ at EC2. *B. avium* strains were electroporated using BIORAD Gene Pulser™ & PULSE CONTROLLER at 2.5 kV, 25 μF, and 400 Ω. All electroporated cells were immediately incubated in SOB with Catabolite repression (SOC) for 1 hour, before selection on antibiotic plates.

### RNA-Isolation

WT *B. avium* and mutant 9b-6 were grown in CDM overnight before being transferred into CDM or TB and allowed to grow overnight. Then, 10^7^ – 10^9^ bacterial cells were lysed in in 1 mL TRI Reagent and incubated at room temperature for 10 minutes. Then, 0.1 mL BCP was added to the homogenate and vigorously shaken for 15 seconds before being incubated at room temperature for 15 minutes. The homogenate was later centrifuged at 12,000 g for 15 minutes at 4°C. Resulting aqueous phase was transferred to a clean Eppendorf tube without disturbing sediment. The sample was DNAse treated by addition of 2U DNAse, 2 μL 10X reaction buffer, and allowed to incubate at 37°C for 45 minutes. Proceeding incubation, the DNAse was deactivated through addition of 2 μL 50 mM EDTA and incubated at 65°C for 30 minutes. Then, 1 μL glycol blue was added to DNAse treated samples for easier visualization of RNA pellet post precipitation. Then, 0.5 mL isopropanol was added to the samples and incubated at room temperature for 10 minutes, before centrifugation at 12,000 g for 8 minutes at 4°C. Resulting RNA pellets were twice cleaned in 1 mL 75% ethanol through centrifugation at 12,000g for 5 minutes at 4°C. The ethanol was then removed and resulting RNA pellet was allowed to dry for 5 minutes before being dissolved in ≤ 30 μL RNAse free water. Resulting DNA and RNA concentrations were measured using QubitTM before being stored at -80°C. RNA was isolated three times per condition, WT *B. avium* in CDM, WT *B. avium* in TB, *B. avium* 9b-6 in CDM, *B. avium* 9b-6 in TB.

### RNA sequencing and transcriptomics

RNA sequencing was conducted by Novogene using NovaSeq PE150. Transcriptome analysis was conducted using the software program Geneious Prime. Each replicate from the respective conditions were assembled to our *B. avium* reference genome (see above). Expression levels were measured for each replicate in their respective conditions using the built-in tool (Annotate & Predict). Differential expression between conditions were measured on the *B. avium* reference genome using the same tool (Annotate & Predict), and DEseq2 was used to normalize raw read counts, and calculate p-values and Log2 ratios (fold change) for each gene.

### GO:term analysis

Up- and down-regulated DEGs, Log2FC ≥ 1 and Log2FC ≤ 1, where isolated from *B. avium* IPDH 591-77 strain-specific GenBank file and translated using a custom python3 script, to ultimately end up as a single FASTA file. FASTA files were uploaded to the software, BioBam (OmicsBox). Amino acid sequenced were blasted using NCBI through OmicsBox, and further analyzed for GO:term enrichment using GO:Slim through OmicsBox (45)

### KEGG Ghost KOALA

Pathway reconstruction of unique and shared DEGs where assembled and analyzed using KEGG Ghost KOALA (46).

## Statistics and reproducibility

All experiments were conducted in experimental triplicate, with statistical analyses being performed using GraphPad Prism (version 10.2.0 (335)).

## Acknowledgments

We would like to thank Dr. Tuan Tran, Dr. Louise Temple-Rosebrook and Dr. Kyle Malter for their advice and discussion on this project. Additionally, we thank Dr. Tom Huxford for providing us with malonate, as well as Dr. Anca Segall for designing and providing the pANT4-pRha-GFP plasmid. Thanks to Hamza Hajama for writing the initial Python script to extract specific data from GenBank files. This work was supported by NIH grant R35 GM146836 and NSF IOS CAREER grant 2143718 to R.J.L., and Harold & June Grant Memorial Scholarship, and Paul G. and Margaret Peninger Memorial Scholarship to N.G.P.

## Supplemental Data

**Supplemental Figure 1.**
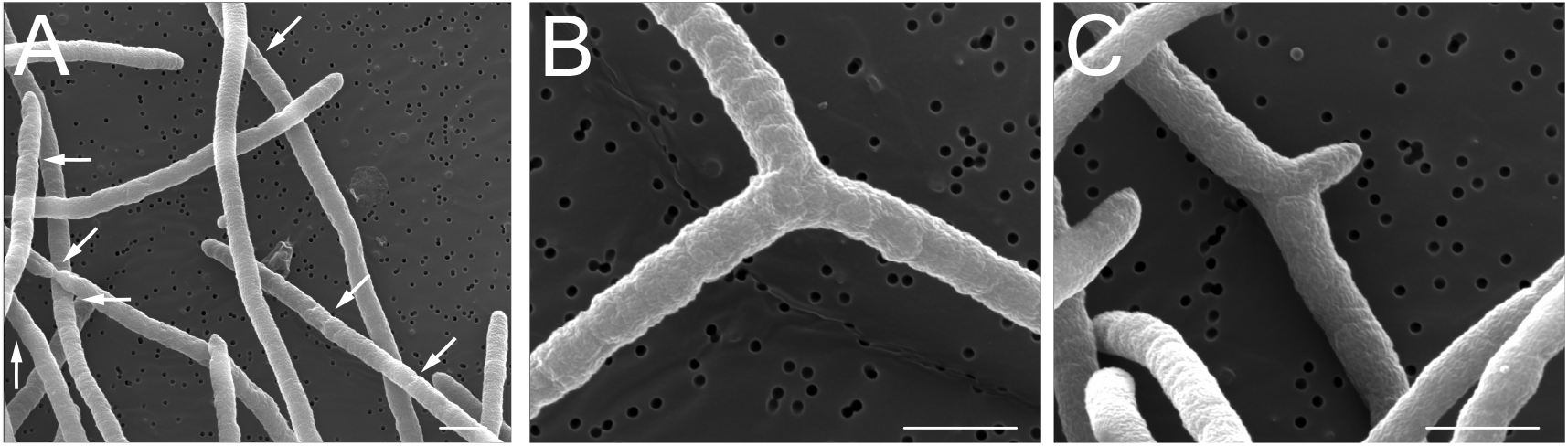
A) Filamentous *B. avium* grown in TB showing early signs of septation indicated by white arrows. B-C) Filamentous *B. avium* grown in TB showing branching. Scale bar=1 µm.

**Supplemental Figure 2.**
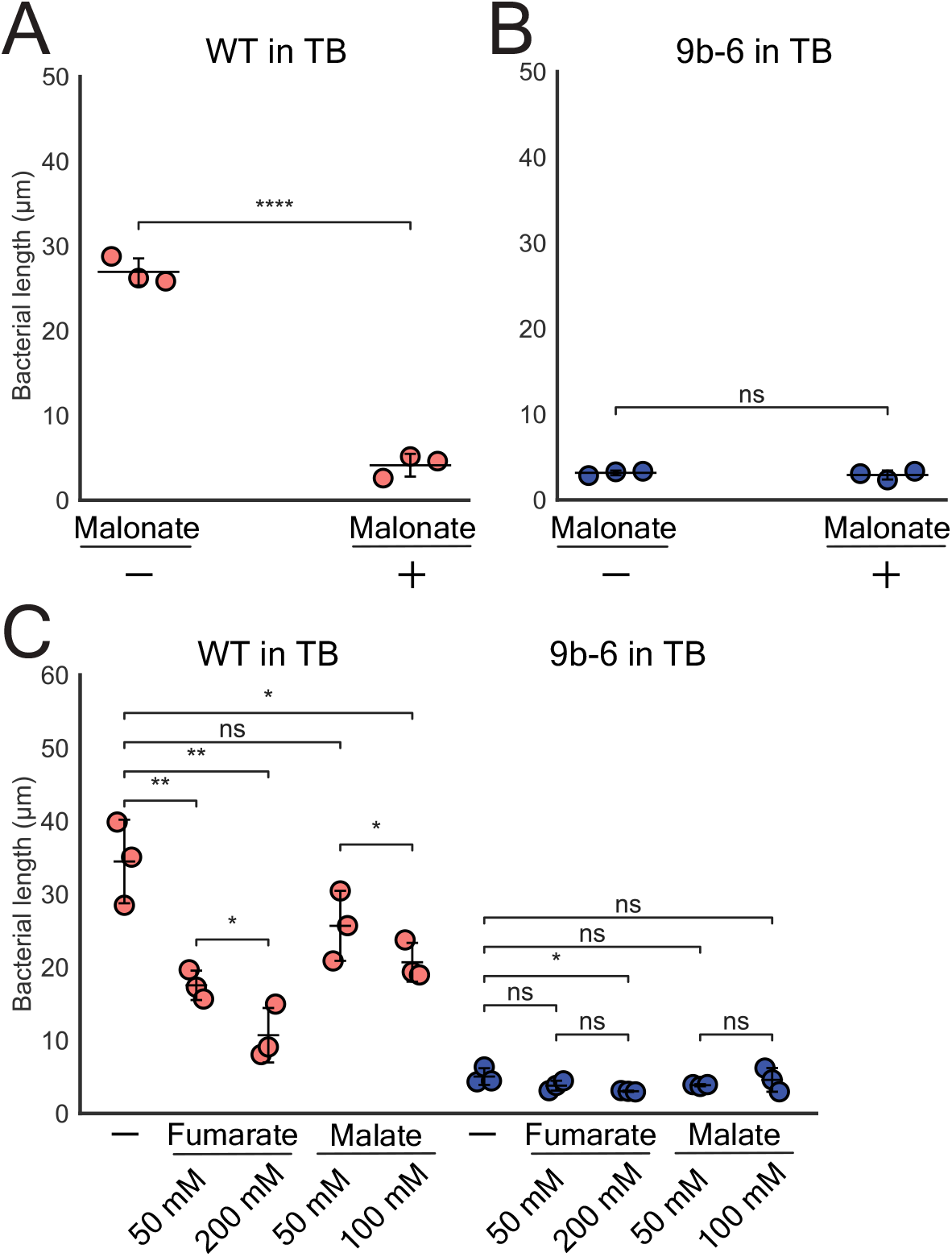
A) WT *B. avium* grown in TB with supplemented malonate (80 mM) showing inhibition of filamentation. B) Mutant 9b-6 grown in TB with supplemented malonate (80 mM) showing no change in bacterial length. C) WT *B. avium* in red and mutant 9b-6 in blue, supplemented with fumarate (50, 200 mM) or malate (50, 100 mM). Experiments were done in triplicate (N=3), n=200/replicate, p<0.05 (*), p<0.01 (**), p<0.0001 (****), ns=not significant, by unpaired two-tailed t-test. Error bars represent SD.

